# Spatially mismatched cerebral blood flow and neuronal activity by intracortical microstimulation

**DOI:** 10.1101/2024.11.10.622852

**Authors:** Alexandra Katherine Isis Yonza, Lechan Tao, Krzysztof Kucharz, Barbara Lind, Xiao Zhang, Kayeon Kim, Changsi Cai

## Abstract

Intracortial microstimulation (ICMS) is widely used in brain-machine interface for neuroprosthetics, particularly with aims of restoring lost sensory and motor functions. However, it remains poorly understood whether neuronal and blood flow responses by ICMS are spatially and temporally matched, as well as their underlying mechanism. Neurovascular coupling (NVC) is the process by which neuronal activity regulates blood flow in the brain to meet local metabolic demands. A hypothetically suboptimal NVC by ICMS may exacerbate the neuronal survival near electrode, contributing to neurodegeneration. In this study, we used wide-field imaging, laser speckle imaging and two-photon imaging in transgenic mice expressing calcium fluorescent indicators in neurons or vascular mural cells to examine this hypothesis. Our results showed blood flow responses are delayed at peak time and prolonged in duration compared with neuronal responses. By varying the stimulation amplitudes, we found that low stimulation intensity lower than 50μA preserved NVC. In contrast, high intensity stimulation caused spatially mismatched NVC, i.e. in the adjacent range within 200∼400 μm from electrode tip, elicited blood flow compromises but neuronal activities increase. Ca^2+^ is the key regulator of contractile tone of vascular mural cells. Our results showed that Ca^2+^ sensitivity at vascular mural cells, i.e. vessel diameter change per unit of Ca^2+^ change, was decreased in the adjacent region of electrode, which partially explained the compromised and mismatched NVC, and which likely further lead to ischemia and neurodegeneration. This study offers a new insight into ICMS-associated neuronal and vascular physiology, and provides an important implication towards optimal design of ICMS: low intensities is more neuroprotective than high intensities by preserving NVC and preventing ischemia. Our discoveries pave the way for new research consideration and contribute to the development of more advanced brain-machine interface.

## Introduction

Artificial neurostimulation via the implantation and direct interface of microelectrodes with the central nervous system has garnered great interest for the chronic mitigation of symptoms derived from various neuropathologies and injuries, as well as the restoration of lost sensory and motor functions [1-5]. Pioneering work with intracortical prosthetics has allowed for advancements to be made in the field of visual restoration, where the dynamic, artificial elicitation of phosphenes has restored a basic semblance of sight to patients blinded by injury and/or disease unrelated to the visual cortex [6, 7].

Despite remarkable results in preclinical studies, clinical and market approvals of intracortical microstimulation (ICMS), studies reported local neuronal restructure and neurodegeneration in adjacent to implanted electrodes [8, 9], causing desensitization and inexplicably large placebo effects following chronic implantation and microstimulation [10]. Besides the non-biological explanation related to device micromotion and suboptimal biocompatibility, many studies have intensively examined and suggested that glial reactivation and neuroinflammation are the main cause of neuronal desensitization and degeneration. At particular, it is attributed to activation, mitigation and proliferation of microglia, astrocytes and oligodenstrocytes [11, 12], as well as the disruption of blood-brain barrier and pericyte deficiencies at capillaries in proximity of implantation site [13, 14]. This forms gliosis, recruits immune cells, promotes inflammation and drives nearby neurons towards excitotoxicity and degeneration.

Notwithstanding these efforts, fewer studies have characterized the neurovascular coupling by ICMS, for which potential decoupling could be a risk factor for neuronal survival. Neurovascular coupling refers to rises in brain activity elicit a rapid increase in local blood flow, providing energy to meet with the local demand. Previous studies have demonstrated neurovascular coupling in ICMS responses [15, 16]. Particularly, increased stimulation intensity leaded to concurrent higher neuronal Ca^2+^ activities and larger hemodynamic responses, and the hemodynamic responses showed a longer latency and duration than neuronal Ca^2+^ [16].

However, the spatial and temporal responses of ICMS-induced neurovascular coupling is yet to be fully understood. And it is crucial to consider whether different subregions of cortex by ICMS demonstrate matched metabolic needs and supplies.

Vascular mural cells are the key element and only contractile compartment in neurovascular unit to control the vessel diameters, and thereby cerebral blood flow [17]. Vascular mural cells are consist of contractile smooth muscles cells on arteries and arterioles, and contractile and non-contractile pericytes on capillaries, as well as non-contractile mural cells on venules and veins [18]. Only contractile mural cells can dilate or constrict vessels, being an important mediator of neurovascular coupling. They ensheath arteries and arterioles, and enwrap the initial branches of capillaries from arterioles. Furthermore, intracellular Ca^2+^ level is a key regulator of vascular mural cells contraction and relaxation [19] in both natural state and upon ICMS [20]. Rise in Ca^2+^ leads to myosin light chain kinase phosphorylation and contraction; Reduction in Ca^2+^ causes dephosphorylation of myosin light chain and relaxation. Thererfore, investigation of Ca^2+^ signals in vascular mural cells and its relationship to blood flow and neuronal activities represents a key underlying mechanism of ICMS-related neurovascular coupling.

Advanced imaging modalities are powerful tools to provide novel insights into ICMS-induced brain responses and its underlying mechanism. Here in this study, we used wide-field imaging, laser speckle imaging and two-photon imaging in living mouse brains to study neuronal, regional blood flow, vascular mural cell Ca^2+^ activities, and single vessel diameter change. We compared the temporal and spatial responses of neurons and blood flow to ICMS, and further examined the vascular mural cell Ca^2+^ and co-localized vessel diameter change at different distances from electrode tip, using multiple transgenic mouse lines. Our results indicate spatially mismatched hemodyanics and ischemia by high intensity ICMS, and provide novel insights microstimulation-induced hemodynamic responses, and contribute to the advancement of ICMS strategies for numerous applications including intracortical sensory prostheses.

## Materials and Methods

### Animal Information and Handling

Both male (25–30 g) and female (20–22 g) mice > 8 weeks of age were utilised throughout this project. Five C57BL/6 mice were used for laser speckle imaging (LSI). Four Acta2-GCaMP8.1-mVermilion (Acta2-GCaMP8.1) mice, carrying calcium indicators in contractile vascular mural cells, were used for two-photon experiments. Two Thy1-GCaMP6 mice (N=2 mice, n=6 different locations), carrying neuronal calcium indicators, were used for wide-field imaging of neuronal activity. All procedures were approved by the Danish National Committee on Health Research in accordance with the European Council’s Convention for the Projection of Vertebrate Animals used for experimental and other scientific purposes.

### Animal surgery

Anesthesia was induced with intraperitoneal injections of xylazine (10 mg/kg) and ketamine (100 mg/kg), dissolved in sterilized water, pH 7.4. Anaesthesia was maintained with intraperitoneal injections of ketamine (50 mg/kg) every 22 min or as needed (as assessed by reflex testing/nociceptive stimulation). Prior to the craniotomies, tracheotomies were performed to ventilate the mice with oxygen and air, in order to keep mouse blood gas in physiological state. Craniotomies were performed to expose an acute cranial window over the right visual cortex immediately prior to imaging sessions. Scalps and connective tissues were surgically removed to cleanly expose the skull, whereupon heads were permanently fixated by glueing bone directly to a specialised aluminium bar. An acute cranial window was created by drilling and removal of a 2 mm skull flap positioned over the right visual cortex (+1 mm anteroposterior, +2 mm mediolateral relative to lambda). The cranial window was kept moist with the regular application of artificial cerebrospinal fluid (aCSF) (in mM: KCl 2.8, NaCl 120, Na_2_HPO_4_ 1, MgCl_2_ 0.876, NaHCO_3_ 22, CaCl_2_ 1.45, glucose 2.55; at 37ºC, aerated with 95% air/5% CO2 to pH 7.4). No mice were recovered following acute surgery, and all mice were humanely euthanized at the end of the imaging and experimentation sessions (3±1 h).

### Protocol of intracortical microstimulations (ICMS)

Microelectrodes (platinum-iridium microelectrodes, PI2PT30.01.A10; Microprobes for Life Sciences) were fixed to a motorized micromanipulator and inserted 200 µm into the exposed cortex at 15–20º. Microelectrodes were connected to ISO-flex (A.M.P.I) units. Cathodic-leading biphasic pulses (200 µs per phase, no interleaving between pulses) were employed in all recordings. All recording sessions utilised a stimulation frequency of 200 Hz. Experiments applied stimulation intensities of 10–200 µA and trial durations of 5 s, with only 1 trial performed per recording. A reference electrode was inserted under the neck skin of the mouse prior to intracortical microelectrode insertion. Counting the highest intensity 200 µA, the charge density is calculated to be lower than safety limit (k = log(Q) + log(Q/A) <1.7), where Q and Q/A represent charge per phase [µC/phase] and charge density per phase [µC cm^-2^/phase], and value k below 1.7 is considered as a safe range [21].

### Wide-field imaging and data analysis

We used the Olympus U-HGLGPS light illumination system and a UPlanSApo 4x objective (0.16 numerical aperture) for wide-field imaging. Image acquisition was at 2Hz with each pixel corresponds to a 1.81µm of tissue, and the field of view was 1440×1920 pixels. To quantify the neuronal response collected from different stimulation sites, we first applied motion correction using two-dimensional (2D) normalized cross-correlation [22], with the first frame as the reference for correction. The region of interest (ROI) corresponding to the stimulated area was then defined by creating a mask, which was based on 3 times standard deviation (SD) above the mean intensity across all pixels in the field of view, following a baseline subtraction and normalization. The frame used for extracting a mask was averaged over the 5-second post-stimulation period. To analyze the neuronal response as a function of distance from the electrode tip, we divided the field into sub-regions, each representing 200µm, arranged as circular vectors radiating from the probe tip. The response amplitude was averaged within each sub-region. To quantify data by pooling across all mice and recording sites, we categorized stimulation intensities into three groups: low (10-50µA) and high (>=50µA). These groups were formed by combining similar stimulation conditions within each range.

### Laser speckle imaging (LSI) and data analysis

The LSI laser source was generated using a VHG-stabilised laser diode (LP785-SAV50, Thorlab) powered by a controller (CLD1011LP, Thorlab). A CMOS camera (acA2040-90umNIR, Basler) and a ×5 0.15-NA objective (HCX PL FLUOTAR, Leica) were utilised for recording. Images were acquired with pixel resolution of 3.19 μm/pixel and image size of 1024×1024 at 30 frames per second (30 Hz). The recording hardware and software was customized as pervious describe [23]. The laser speckle images were post-processed using Matlab 2023a with custom-made code [24]. Absolute blood flow index (absBFI) was calculated by inverted square of laser speckle contrast intensity at each pixel. Every 15 frames (half a second duration) were averaged to represent absBFI at each consecutive half second. Baseline absBFI was calculated by averaging all frames in a 5s time period before the onset of stimulation.

Relative blood flow index (relBFI) was calculated as follow:

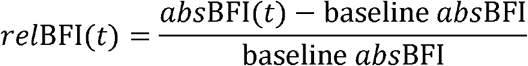

Attenuated blood flow change at the electrode tip was calculated as follow:

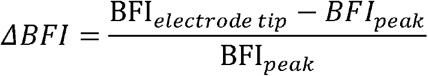

### Two-photon imaging and data analysis

The FluoView FVMPE-RS (Olympus) two-photon microscope equipped with a 25□×□1.05 NA water-immersion objective, the Mai-Tai DeepSee femtosecond laser, and GaAsP detectors was employed for experiments and recording under TPM. An excitation wavelength of 920 nm was utilised for Acta2-GCaMP8.1 mouse recordings. 40 uL Texas Red was injected intravenously via retro-orbital injection. Image acquisition frequency remained around approximately 1.8 Hz across all recordings. A custom motion correction algorithm was applied to TPM recordings. A small region background region where pixel fluorescence remains stable and dark over the course of the recording was selected as the reference for background-subtraction from all frames. ROIs were manually drawn to include fluorescent structures of interest as well as a portion of surrounding background. Background vs. foreground was distinguished by a threshold value calculated as mean pixel intensity + 2*STD so as to correctly identify and encompass foreground structures. The frame containing the onset of stimulation was denoted as the baseline to normalise subsequent stimulation trial frames to. ΔF/F_baseline_ was plotted over time of given stimulation trials for each ROI drawn.

### Statistics

We performed unpaired t-test to compare the response between different intensities, and region of interest defined as a function of distance from stimulating electrode (p<0.05). To assess the relationship between neuronal amplitude distance from the electrode tip, we used Pearson’s correlation with statistical significance at p<0.05.

## Results

### Neuronal response decreases with distance from electrode tip

To define the spatial profile of neuronal responses to ICMS, we used Thy1-GCaMP6 mice expressing green florescent calcium indicator in neurons, and investigated neuronal responses to ICMS at mouse primary visual cortex (Fig. 1A). We used wide-field imaging in anesthetized mice to examine the activated brain region by ICMS. Similar with the previous studies [16], activated area of neuronal calcium responses increase with elevated stimulation intensities (Fig. 1B), where the frames within first 5 seconds post-stimulation were averaged. The time course and peak amplitude of neuronal responses in the whole field of view also increases by rising stimulating intensity (Fig. 1C). Next, we asked whether neuronal response amplitudes decrease with larger distance from electrode tip. We therefore assigned the concentric rings surrounding electrode tip with inter-ring distance of 200 μm (Fig. 1D left), all the pixel intensity from the each circular ring area are averaged to represent the response intensity at this distance from electrode tip. As expected, the response intensity varies as a function of distance to electrode tip with highest neuronal response closest to the tip. The same phenomena were observed across all the stimulation intensities (Fig. 1D right). After categorizing stimulation intensities into two groups: low (<50µA) and high (>=50µA), our population level analyses confirmed that response peak amplitudes significantly increased with higher stimulation intensities (Fig. 1E, paired t-test, p<0.05). Additionally, the response amplitudes significantly decreased as the distance from the electrode gets farther across all stimulation intensities (Fig. 1F).

**Figure 1.**
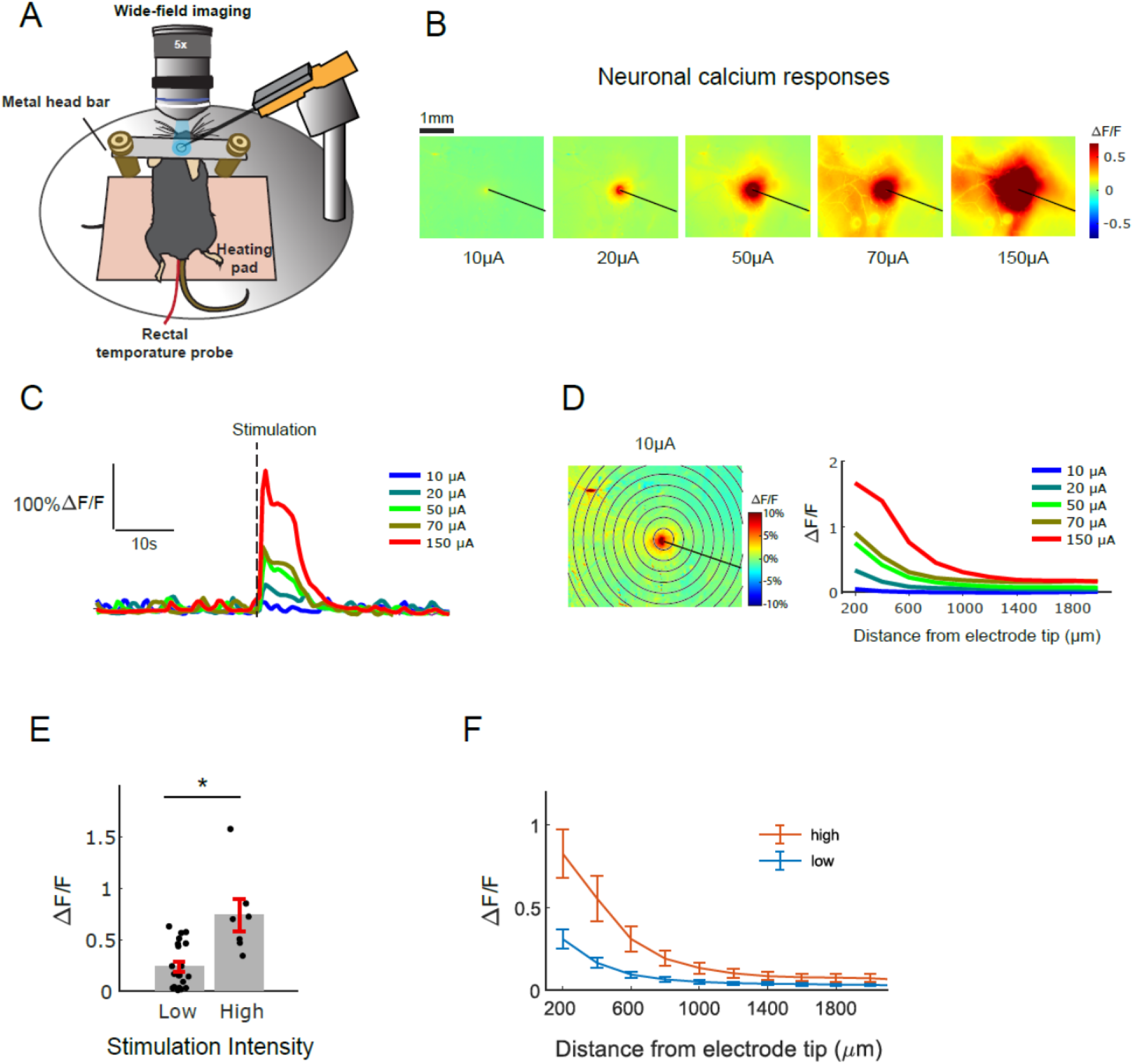
Neural responses to ICMS under wide-field imaging. **(A)** The experimental diagram of in vivo wide-field imaging. **(B)** An example recording site showing a baseline corrected frame, averaged over a 5-second period following stimulation, during 10∼150µA stimulation intensities. The black line indicates the electrode of the location. **(C)** The ΔF/F time course from panel B is plotted, with the gray bar indicating the stimulation period. **(D)** Left: An example response during 10µA stimulation, showing the circular vector rings with 200µm sub-regions. Right: The response amplitude is plotted as a function of distance from the electrode tip (in µm). **(E)** Pooled data from all mice and recording/stimulation sites. Each dot represents ΔF/F averaged over the 5-second post stimulation period from a single trial, plotted against stimulation intensity categories. **(F)** Same convention as E showing population response, but plotted response amplitude (ΔF/F) as a function of distance from electrode tip, during low to high stimulation intensities. Low intensities <50μA; High intensities > 50μA. Data are given as the mean ± s.e.m. *p < 0.05, **p < 0.01, ***p < 0.001

### Blood flow response mismatches with neuronal response in the field of view

In natural stimulation, higher neuronal activities are co-localized with higher blood flow responses, in order to match the metabolic need and delivery [25]. We examined whether ICMS-elicited blood flow responses are in the same temporal and spatial pattern with neuronal responses. In these experiments, we used laser speckle imaging at primary visual cortex of anesthetized wild-type mice by ICMS (Fig. 2A). A representative time course of absolute blood flow index (absBFI, Fig. 2B upper row) and relative blood flow index (relBFI, Fig. 2B lower row) by 70μA electrical stimulation showed blood flow increase in the adjacent region of electrode tip. Similar with neuronal responses, the time course and peak amplitudes of blood flow responses increased with rising stimulation intensities (Fig. 2C). Interestingly, the peak time and duration of blood flow responses are prolonged comparing with neuronal responses (Fig. 2D-E). Blood flow responses peaked at 3s post stimulation, in comparison neuronal responses peaked at 1∼2s. Further, we examined whether the blood flow responses are spatially matched with neuronal responses, i.e. decrease with distance to electrode tip. Surprisingly, we found that the blood flow was peaked away from the tip, using the same definition of concentric rings (Fig. 2F-G).

**Figure 2.**
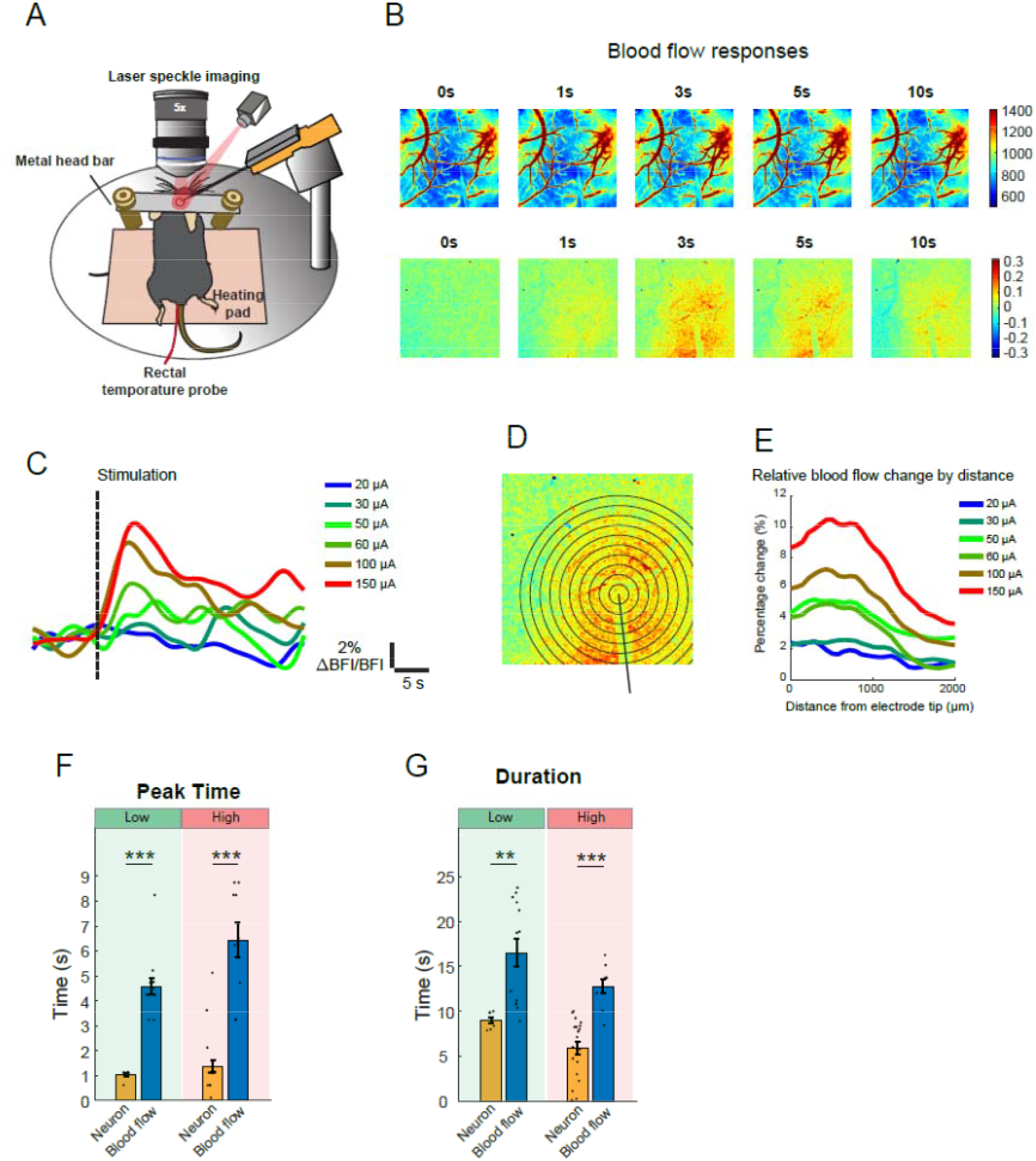
Blood flow responses to ICMS under laser speckle imaging. **(A)** The experimental diagram of in vivo laser speckle imaging. **(B)** Image cascade of absolute blood flow index (upper row) and relative blood flow index (lower row). **(C)** Time course of relative blood flow change by different stimulation intensities. **(D)** Representative image of concentric rings to calculate blood flow change as a function of distance to electrode tip. **(E)** Overlay of curves as distance-related blood flow change under different stimulation intensities. **(F)** Comparison of peak time of neuronal responses and blood flow responses at both low and high stimulation intensity. **(G)** Comparison of duration of neuronal responses and blood flow responses at both low and high stimulation intensity. Data are given as the mean ± s.e.m. *p < 0.05, **p < 0.01, ***p < 0.001

### Blood flow response in parenchyma peaks away from electrode tip

Laser speckle imaging detects the deflection of laser by the moving particles in the blood vessels, and thereby LSI signal intensity is much higher in arteries and arterioles than capillaries in parenchyma [26] (Fig. 3A). If considering the whole field of view, the blood flow responses may artificially peak at the location of big pial arteries. Further, parenchymal blood flow response, i.e. capillary flow change, reflects local blood supply to the regional activated and starving neurons. Therefore, we decide to mask away all the big pial vessels and the shadow of microelectrode, and to only consider the parenchymal blood flow change (Fig.3B). By increasing amplitudes of ICMS from 20μA to 200μA, parenchymal blood flow was elevated in the field of view, with averaging across the initial 5 seconds post stimulation (Fig. 3C). And the area in adjacent to the electrode tip showed dampened blood flow change compared with region away from the tip. We further quantified the relative blood flow change as a function of distance from the tip at different stimulation intensities (Fig. 3D). Attenuated blood flow change at the electrode tip was calculated based on the equation in the Methods section, and the results showed linear increase with higher stimulation intensity (Fig. 3E). This represents a more severe mismatch between blood flow demand and delivery at higher stimulation intensities. Furthermore, the location of response peak moved further away from the electrode tip with increased intensity (Fig. 3F). Although the peak amplitude of the regional flow responses showed a linear rise with higher intensity (Fig. 3H), the blood flow response at the electrode tip increased at lower stimulation intensity but reached plateau when stimulation intensity is higher than 70μA (Fig. 3G). Population analysis across all mice suggested similar results (Fig. 3I-L), with significant difference between low and high intensities.

**Figure 3.**
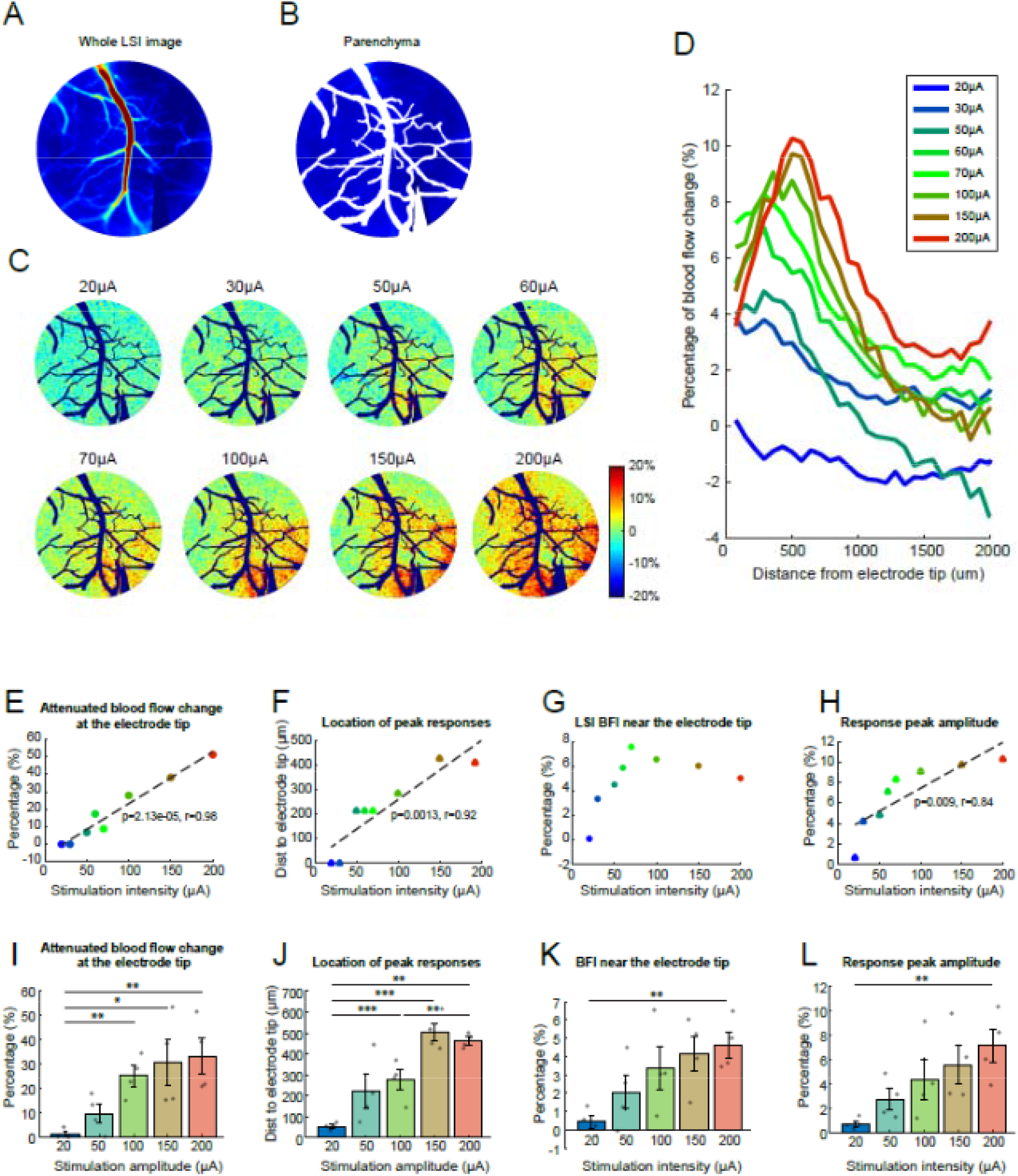
Parenchymal blood flow responses to ICMS under laser speckle imaging (LSI). **(A)** Representative LSI images in the whole field of view. **(B)** By masking away the pial arteries and pial veins, only parenchymal blood flow is considered. **(C)** Representative images of averaged blood flow responses across 0∼5s post-stimulation under different stimulation intensities. **(D)** Based on images in C, the overlay of curves as distance-associated blood flow change under different stimulation intensities. **(E-H)** Based on curves in D, quantifications are plotted as a function of stimulation intensities: (**E**) Attenuated blood flow change at the electrode tip compared with peak blood flow; (**F**) Location of peak responses; (**G**) Blood flow responses at the electrode tip; (**H**) Response peak amplitudes. **(I-L)** Populationary analysis of all mice (N=4, n=4) as the same measurement of (E-H). Data are given as the mean ± s.e.m. *p < 0.05, **p < 0.01, ***p < 0.001

### Ca^2+^ sensitivity underlies spatially different vascular responses to ICMS

Next, we asked what is the possible mechanism for attenuated vascular responses in adjacent to microelectrode tip. Vascular mural cells are the only contractile cell types in neurovascular unit, and are well documented to dominate the vessel diameter changes at arterioles and capillaries. The contractile tone of vascular mural cells are regulated by their intracellular Ca^2+^ level.

Therefore, we used transgenic mice Acta2_GCaMP8 expressing Ca^2+^ fluorescent indicator GCaMP8 in contractile mural cells, and employed in vivo two-photon microscopy to examine the Ca^2+^ change in vascular mural cells in response to ICMS. Further, we injected Texas Red intravenously to stain vessel lumen as red color. Under wide-field imaging modality, we focused on the cortical surface and found vasculature of interest using 5x objective (Fig. 4A), and further switched to 25x objective to carefully pick three different penetrating arterioles (PAs) (Fig. 4B). Upon 70μA intensity of ICMS, vessel dilation was accompanied with Ca^2+^ decrease in vascular mural cells in the representative two-photon imaging cascades (Fig. 4D-F). Based on the distance between each PA and the electrode tip, we classified PAs to near (<200 μm), medium (200∼400 μm) and far (> 400 μm). The waveforms of diameter change and vascular mural cell Ca^2+^ were shown in Fig. 4G-I at each location. The peak amplitude and area under curve (AUC) of vessel diameter change was largest at the medium distance region; while peak amplitude and AUC of mural cell Ca^2+^ were largest at the near distance region (Fig. 4J&L).

**Figure 4.**
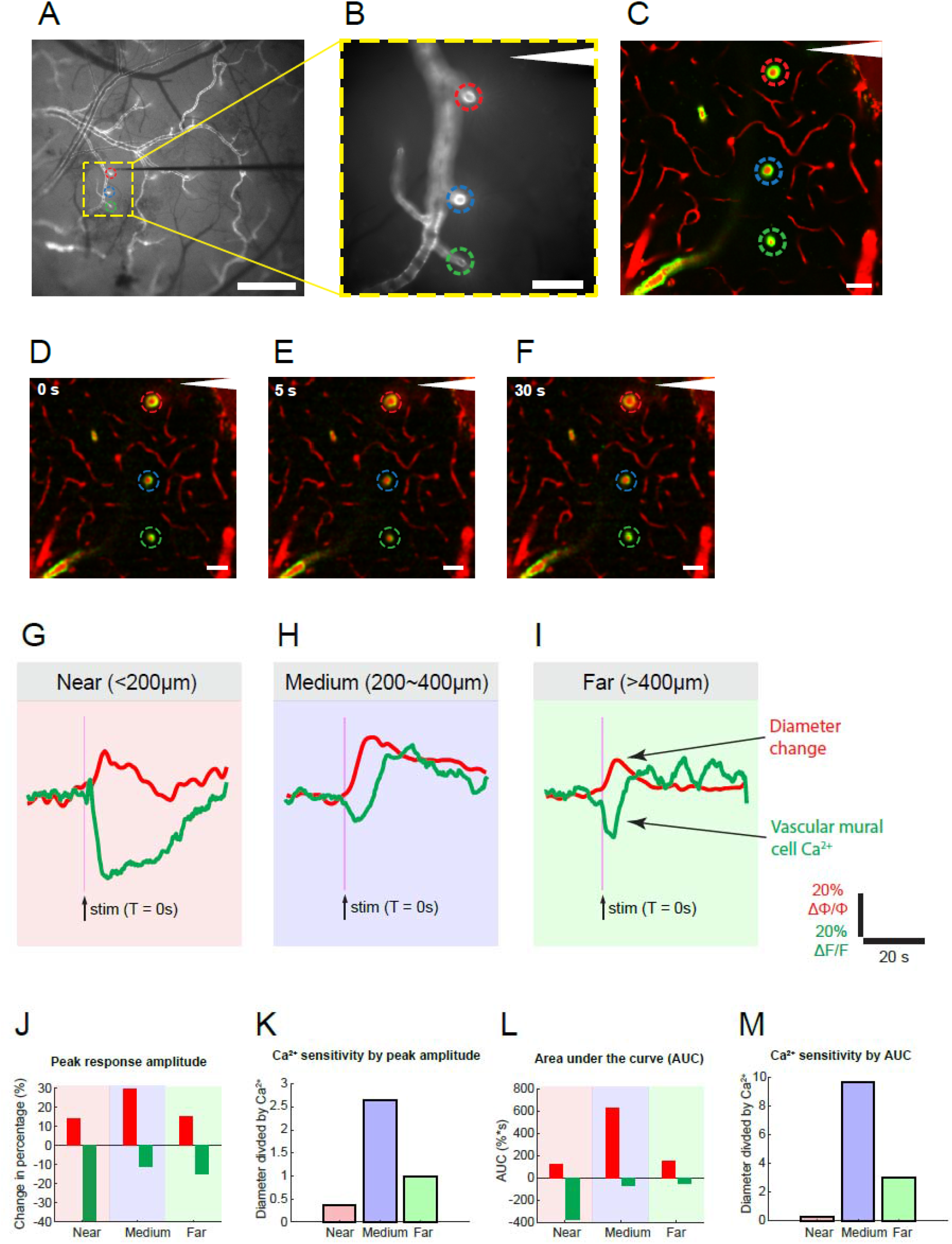
In vivo two-photon imaging of vessel diameter and Ca^2+^ signals in Acta2_GCaMP8 mice in response to ICMS. **(A)** Representative wide-field imaging of cortical surface of a Acta2_GCaMP8 mouse with an inserted microelectrode in primary visual cortex. The image was taken under 5x objective. Scale bar 500 μm. **(B)** After switching to 25x objective, the wide-field imaging of the same vasculature as (A). Scale bar 100 μm **(C)** Two-photon imaging of the same vasculature. Green: green fluorescent calcium indicators in contractile mural cells with promoter of Acta2. Red: Texas Red injected intravenously to stain vessel lumen. Scale bar 50 μm. **(D-F)** Two-photon imaging cascades of vessel diameter change and vascular mural cell Ca^2+^ upon 70 μA ICMS. Three penetrating arterioles are in the same color-code with (A-C). **(G-I)** The paired curves of diameter change and co-localized vascular mural cell Ca^2+^ at the three locations. **(J)** Comparison of response peak amplitudes of vessel diameter and Ca^2+^. **(K)** Comparison of Ca^2+^ sensitivity at the three locations, using vessel amplitude divided by Ca^2+^ amplitude. **(L)** Area under the curve is calculated and compared, similar with (J). **(M)** Ca^2+^ sensitivity is calculated and compared based on AUC.

Considering vascular mural cell Ca^2+^ is a key determinant of vessel diameter, we defined the Ca^2+^ sensitivity as dividing diameter change by Ca^2+^ change at each vascular location. Vascular mural cell Ca^2+^ showed highest sensitivity at the medium distance region by calculating both peak amplitudes (Fig. 4K) or AUC (Fig. 4M), indicating that smaller Ca^2+^ changes drives larger diameter changes in this region. Populational analysis including all mice showed insignificant but trendy larger vasodilation at medium region, but with significant Ca^2+^ drop at near region, especially calculated by AUC (Fig. 5A-D). Collective comparison of Ca^2+^ sensitivity showed significant higher at medium region than other two regions (Fig. E-F).

**Figure 5.**
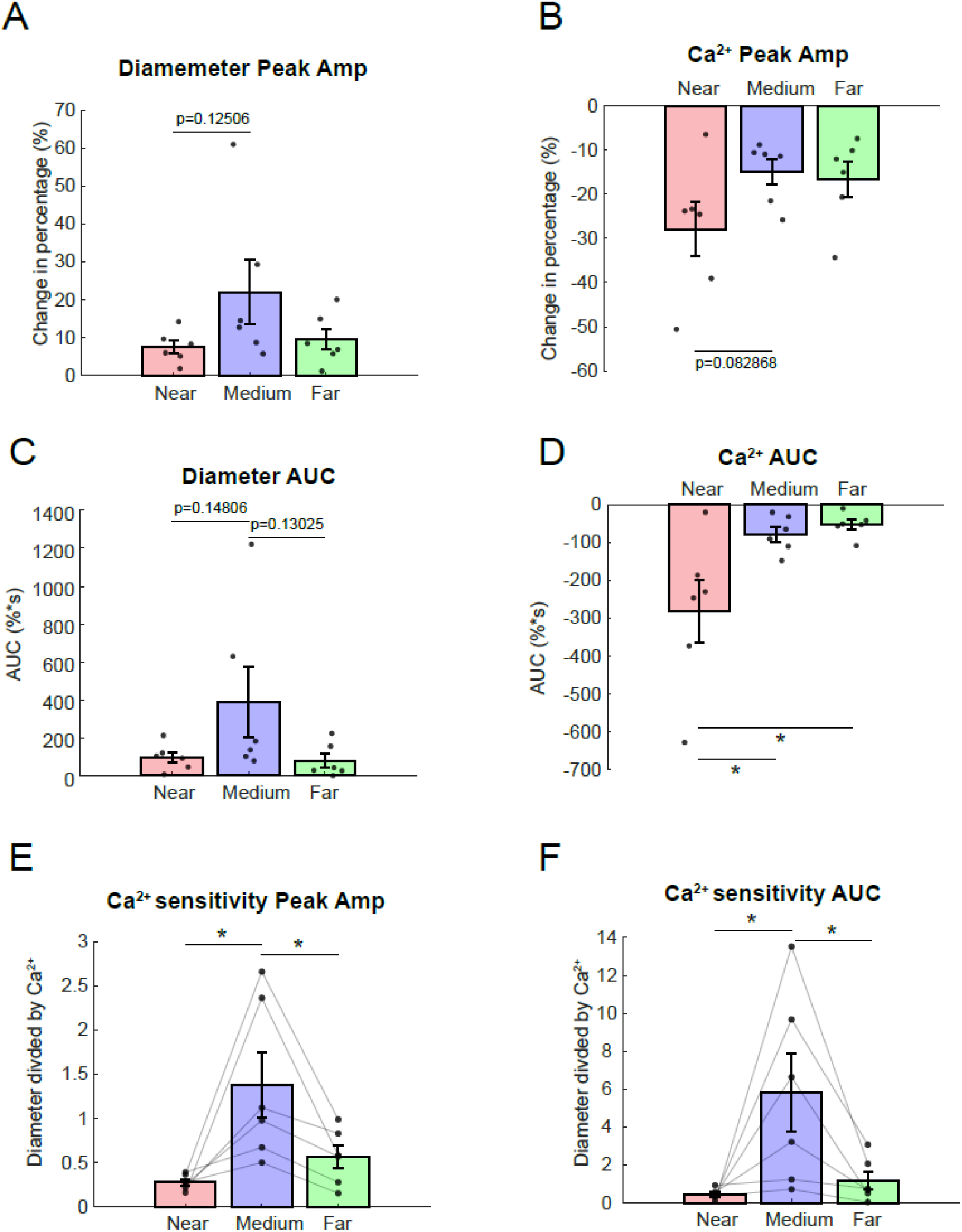
Collective analysis of Ca^2+^ sensitivity upon high stimulation intensity. The following measurements are compared: **(A)** Response peak amplitude of diameter changes. **(B)** Response peak amplitude of vascular mural cell Ca^2+^. **(C)** Response area under the curve (AUC) of diameter changes. **(D)** Response AUC of vascular mural cell Ca^2+^. **(E)** Ca^2+^ sensitivity calculated by peak amplitude. **(F)** Ca^2+^ sensitivity calculated by AUC. Data are given as the mean ± s.e.m. *p < 0.05, **p < 0.01, ***p < 0.001

## Discussion

In this study, we showed that ICMS induced activation in neuronal Ca^2+^ signals and elevated blood flow responses in visual cortex of anesthetized mice. The neuronal response amplitude increased with higher stimulation intensity, and the response amplitude decreased with larger distance from the electrode tip. In contrast to neuronal responses, blood flow responses were delayed, prolonged, and their spatial pattern differed between low and high stimulation intensities. Low stimulation intensity elicited similar spatial pattern in neuronal and blood flow responses. High stimulation intensity attenuated the blood flow responses in adjacent to electrode, and the spatial extension of this attenuation was dependent on stimulation intensity. Furthermore, the attenuation of blood flow responses near electrode is partially attributed to the decreased Ca^2+^ sensitivities in contractile vascular mural cells in adjacent to electrode compared with distal region. This indicates a metabolic mismatch and shortage between demand and delivery closely surrounding the electrode tip, leading to ischemia, which unveil a new mechanism contributing to the neurodegeneration upon ICMS. One important implication is that low intensity stimulation preserves neurovascular coupling better than high intensity stimulation, and therefore potentially benefits neuronal survival upon implantation and stimulation. Overall, our results provide valuable insights into ICMS-associated neuronal and vascular physiology, and inspire the further development of biomimetric prostheses.

### Possible mechanism of varied Ca^2+^ sensitivities in vascular mural cells

Vascular mural cells are categorized in different subtypes, and their genes, morphologies, localizations and physiological functions vary in a wide spectrum [27]. Here in this study, our focus was ICMS-related hemodynamic responses and neurovascular coupling, and thereby only contractile vascular mural cells were considered, which include arterial smooth muscle cells and contractile pericytes at the initial capillary branches from arterioles [28, 29]. Intracellular Ca^2+^ is the key molecule determining contractile tone of the vascular mural cells. Therefore, we used two-photon microscopy to measure synchronically Ca^2+^ signals in vascular mural cells and vessel diameter change in Acta2_GCaMP8 mice, where Acta2 is specific promoter for contractile mural cells. Their Ca^2+^ sensitivity refer as the amount of vessel diameter change induced by per unit Ca^2+^ change. Previous vascular studies reported that in pathological conditions such as aging or brain ischemic stroke, Ca^2+^ sensitivity increases [30]; while other diseases showed decreased Ca^2+^ sensitivity [31]. In this study, we reported decreased Ca^2+^ sensitivity in contractile vascular mural cells in adjacent to electrode tip, which may underlie the compromised blood supply in the local region. A few possible molelcular and cellular pathways could be further considered: (1) In physiological condition, RhoA/Rho-Kinase signaling modulates Ca^2+^ sensitivity in vascular mural cells. High intensity electrical stimulation may impair RhoA/Rho-Kinase signaling and lead to desensitization of Ca^2+^ in contractile vascular mural cells near the electrode. (2) Acute insertion of microelectrode causes blood-brain barrier break down, microglia migration and elevated level of proinflammation cytokines [12]. This may disrupt actin-myosin interaction and myosin light chain kinase in contractile vascular mural cells, and therefore compromise the relaxation. (3) Contraction and relaxation of vascular mural cells could also be mediated by Ca^2+^-independent pathway, for example via intracellular secondary messengers cyclic guanosine monophosphate (cGMP) and cyclic adenosine monophosphate (cAMP) [32]. Particularly, cGMP is attributed to nitric oxide (NO) which is synthesized in neighboring endothelial cells. Disruption of endothelial cells by electrode insertion and stimulation may lead to decreasing synthesis of NO, weakening its contribution to neurovascular coupling, and thereby attenuating blood flow responses in adjacent region of electrode compared to distal region. These proposed mechanisms should be examined in the future studies.

### Spatially mismatched neurovascular coupling by high intensity stimulation

Our study reported a potentially important result: By low intensity microstimulation (threshold level), neurovascular coupling is spatially preserved, i.e. high/low neuronal response is co-localized with high/low blood flow responses, respectively, as a function of distance to electrode tip. However, by high intensity microstimulation, neurovascular coupling is spatially mismatched in 200 ∼ 400 μm radius region from the electrode. The closer distance to electrode, the larger mismatch is taken place. This likely leads to ischemia, and contributes to shortage of metabolic supply and neurodegeneration in adjacent region of electrodes. Actually, the neurovascular decoupling has been proposed to be the major cause of some brain diseases. For example, ischemic stroke where blood flow is blocked. The starving neurons by insufficient blood supply in penumbra region of stroke demonstrate progressive apoptosis and lead to enlarged infarcted brain region over time [33]. Another example is brain vascular aging which is associated with excessive stiffness and thickness of vessel wall, and give rise to compromised blood flow upon neuronal activation. This is associated with cognitive decline and dementia [32]. Therefore, keeping intact and well-functioning neurovascular coupling by ICMS is previously overlooked but potentially important topic. In light of our results, we recommend to use threshold-level low intensity stimulation.

On the other hand, the temporal factors of neurovascular coupling is likely to be preserved by both low and high intensity ICMS. Our results showed that the blood flow responses by ICMS is delayed in peak and prolonged in duration than neuronal responses. This is in accord with the previous studies using intrinsic optical imaging and two-photon microscopy to examine the neurovascular coupling by physiological stimulation [34] and ICMS [16]. Although prolonged blood flow responses may potentially benefit the survival of neurons surrounding electrode even if there is spatial mismatch during stimulation, there are nonmetabolic functions of neurovascular coupling [35]. The spatial mismatch of nonmetabolic functions may not be compensatable by prolonged temporal responses, for example, secretion and circulation of cerebrospinal fluid [36] heat removal [37] and feedback alteration of neighboring neuronal excitability [38].

### Limitations and future directions

Despite previous intensive studies on neurophysiology by brain stimulations employed anesthetic animals as in vivo model in the past few decades, it is acknowledged that the excitability of neurons and blood flow responses are significantly decreased by anesthestisia [39]. In this study, we used ketamine in all neurovascular coupling related experiments. Ketamine is an NMDA receptor antagonist, which disrupts neuronal excitatory signaling. Although ketamine was reported to preserve neurovascular coupling better than other anesthesia [40], it is still possible that ICMS-associated neurovascular coupling may differ in anesthetic state from awake state. Therefore, re-examination of our results in awake mice will be valuable.

Furthermorein this study, we examined the neuronal and blood flow responses in acutely prepared mice. However, brain response to acute and chronic implantation and microstimulation may differ. A few progressive alteration related to neurovascular coupling should be taken into consideration: (1) Upon acute insertion, microglia are activated and migrate to the injury site [12]. Microglia are recently reported to actively modulate neurovascular coupling [41, 42]; (2) Within days, astrocytes become reactive and proliferated, forming glial encapsulation surrounding the electrode, and further promote inflammation. Astrocytes are known to regulate neurovascular coupling via their endfeet closely enwrapping of and communicating with vascular mural cells [43]. (3) Vascular endothelial cells are recently suggested to be ‘vascular information high way’ to conduct fast propagating electrical signals, conveying the metabolic needs from capillaries to pial arteries [44]. Disruption of blood-brain barrier and impairment of endothelial cells by electrode insertion and chronic inflammation likely compromise neurovascular coupling. (4) Pericytes constrict in acute phase of implantation and leads to vasoconstriction. And they further proliferate and facilitate angiogenesis in chronic term [20]. The density, morphology and functional alteration of pericytes may reshape the neurovascular coupling after chronic implantation. Thereby, re-examination of our results in a longitudinal study will be the next step.

## Conclusion

In this study, we examined an important question: whether intracortical microstimulation elicits temporally and spatially matched neurovascular coupling. Using multimodal imaging on transgenic mice, we found that blood flow response is temporally delayed and prolonged comparing with neuronal responses. However, the blood flow response is spatially mismatched by high intensity stimulation, i.e. blood flow is attenuated within the region of 400μm radius from electrode tip, compared with peaked neuronal activities in this region. This exacerbates the metabolic deficit and potentially leads to neurodegeneration. This phenomena is attributed to the decreased calcium sensitivity at contractile vascular mural cells in adjacent to electrode.

However, if delivering low intensity stimulation, neurovascular coupling is spatially matches. Our study provides important implication for the stimulation parameters of intracortical prosthetic device – low intensity stimulation preserves neurovascular coupling in adjacent area of electrode, and therefore are more neuroprotective than high intensity.

## Acknowledgements

This study was supported by the Lundbeck Foundation, the Danish Medical Research Council, the Novo Nordisk foundation and a Nordea Foundation Grant to the Center for Healthy Aging, the Alice Brenaa Foundation, Augustinus Foundation, Carl og Ellen Hertz Familielegat, A. P. Møller Foundation, Helsefonden, Dagmar Marshall Fond and Torben og Alice Frimodts Fond.

